# Evidence that the cell cycle is a series of uncoupled, memoryless phases

**DOI:** 10.1101/283614

**Authors:** Hui Xiao Chao, Randy I. Fakhreddin, Hristo K. Shimerov, Rashmi J. Kumar, Gaorav P. Gupta, Jeremy E. Purvis

**Affiliations:** Department of Genetics, University of North Carolina at Chapel Hill, 120 Mason Farm Road, Chapel Hill, NC 27599-7264; Cirriculum for Bioinformatics and Computational Biology, University of North Carolina at Chapel Hill, 120 Mason Farm Road, Chapel Hill, NC 27599-7264; Curriculum in Genetics and Molecular Biology, University of North Carolina at Chapel Hill, 120 Mason Farm Road, Chapel Hill, NC 27599-7264; Lineberger Comprehensive Cancer Center, University of North Carolina at Chapel Hill, 120 Mason Farm Road, Chapel Hill, NC 27599-7264; Department of Radiation Oncology, University of North Carolina at Chapel Hill, 120 Mason Farm Road, Chapel Hill, NC 27599-7264

## Abstract

The cell cycle is canonically described as a series of 4 phases: G1 (gap phase 1), S (DNA synthesis), G2 (gap phase 2), and M (mitosis). Various models have been proposed to describe the durations of each phase, including a two-state model with fixed S-G2-M duration and random G1 duration**^1,2^**; a “stretched” model in which phase durations are proportional**^3^**; and an inheritance model in which sister cells show correlated phase durations**^2,4^**. A fundamental challenge is to understand the quantitative laws that govern cell-cycle progression and to reconcile the evidence supporting these different models. Here, we used time-lapse fluorescence microscopy to quantify the durations of G1, S, G2, and M phases for thousands of individual cells from three human cell lines. We found no evidence of correlation between any pair of phase durations. Instead, each phase followed an Erlang distribution with a characteristic rate and number of steps. These observations suggest that each cell cycle phase is memoryless with respect to previous phase durations. We challenged this model by perturbing the durations of specific phases through oncogene activation, inhibition of DNA synthesis, reduced temperature, and DNA damage. Phase durations remained uncoupled in individual cells despite large changes in durations in cell populations. To explain this behavior, we propose a mathematical model in which the independence of cell-cycle phase durations arises from a large number of molecular factors that each exerts a minor influence on the rate of cell-cycle progression. The model predicts that it is possible to force correlations between phases by making large perturbations to a single factor that contributes to more than one phase duration, which we confirmed experimentally by inhibiting cyclin-dependent kinase 2 (CDK2). We further report that phases can show coupling under certain dysfunctional states such as in a transformed cell line with defective cell cycle checkpoints. This quantitative model of cell cycle progression explains the paradoxical observation that phase durations are both inherited and independent and suggests how cell cycle progression may be altered in disease states.

---

The discovery that DNA synthesis occurs during a well-defined period of time between cell divisions^5^ led to the development of the canonical four-stage cell cycle model comprising G1, S, G2, and M phases. It has long been known that the durations of these phases can vary considerably across cell types^6^. For example, stem cells and immune cells have relatively brief G1 durations compared to somatic cells^7–9^. Phase durations can also change under certain environmental stresses such as starvation, which lengthens G1^10^, or DNA damage, which prolongs G1 and G2^11,12^. Furthermore, examination of individual cells has revealed that phase durations vary even among clonal cells under similar environmental conditions^6^. These apparently stochastic differences in cell-cycle durations were originally attributed exclusively to the G1 phase^13^. However, more recent studies in multiple cell types have revealed that S and G2 also contribute significant variability to total cell cycle duration^3,14,15^. Collectively, these studies have revealed that differences in cell cycle durations are an inherent property of individual cells and raise the fundamental question of how these durations are determined.

Over the past 50 years, multiple models have been put forth to explain the differences in cell cycle phase durations among individual cells. By measuring the time between consecutive cell divisions in unsynchronized cells, Smith and Martin proposed a probabilistic model in which the cell cycle is composed of a random part (“A-state”) that includes most of G1, and a determinate part (“B-phase”) composed of the combined S-G2-M phases and the remaining duration of G1^1^. The widely accepted implication of this model is that variability in total cell cycle duration stems mostly from G1 and that the durations of the A-state and B-phase are uncorrelated (since one is fixed and the other is random). However, a more recent body of work using time-lapse fluorescence microscopy suggests that cell-cycle phase durations may be correlated. Using the FUCCI fluorescent reporter system^16^ to estimate the onset of S phase in proliferating mouse lymphocytes, the duration of the combined S-G2-M phase was reported to be proportional to the total cell cycle duration^3,4^. This “stretched” cell cycle model suggests that S-G2-M contributes a substantial amount of variation to total cell cycle duration and claims that a shared factor affects progression through multiple phases^3^. Along these lines, Araujo *et al*. recently reported that each phase is correlated with total cell cycle length with the exception of M phase, which is “temporally insulated” from upstream events^17^. However, these claims are based primarily on comparison of individual phase durations to total cell cycle length rather than direct comparison of the durations of each phase.

The possibility of phase coupling is supported by the observation that many biochemical processes are known to exert control over more than one phase. For example, expression of the E2F family of transcription factors, which target genes involved in the G1/S and G2/M transitions and replication, influences the durations of G1, S, and G2^18–22^. Furthermore, certain stress signals, such as those evoked by DNA damage, can be transmitted from one phase to the next and even inherited from a mother cell’s G2 to the daughter’s G1^11,23^. The existence of molecular factors that control phase durations is also consistent with the observation that sister cells show strong correlations in their phase durations^4,24^. Recent quantification of G1 and S-G2-M showed that G1 itself is heritable and highly correlated between sisters^4^. Reconciling the heritable nature of phase durations with the question of phase coupling is necessary for building a comprehensive picture of cell cycle progression in individual cells.

Here, we report precise measurements of G1, S, G2, and M phase durations in three human cell lines. We find that each phase operates according to a distinct timescale and detect no evidence of phase coupling. Instead, phase progression can be accurately modeled as a sequence of memoryless steps in which the duration of each phase is independent of the previous phase duration. This lack of correlation holds even when phase durations are altered by external stresses, although under certain conditions of extreme perturbation or defective checkpoints, phase coupling can be introduced. To explain these observations, we propose a mathematical model in which a large number of heritable factors can each weakly couple the durations of individual phases, but in ensemble the phases are effectively uncoupled. This quantitative description of cell-cycle progression provides a new conceptual framework for studying diseases in which cell cycle progress is dysregulated.

## Results

### Cell cycle phase durations are uncoupled under unstressed conditions

We examined cell cycle progression in three human cell lines: a non-transformed cell line (hTERT RPE-1, abbreviated RPE), a transformed osteosarcoma cell line (U2OS), and an embryonic stem cell line (H9). RPE cells are non-transformed human epithelial cells immortalized with telomerase reverse transcriptase with intact cell cycle regulators^25^; U2OS cells are transformed cancer cell line with near triploidy and an unstable G1 checkpoint^26–28^. H9 cells are derived from human blastocysts^29^ and exhibit rapid proliferation characterized by a shortened G1 duration^7^. We used the proliferating cell nuclear antigen (PCNA)-mCherry fluorescent reporter to quantify, for each cell, the duration of G1, S, G2, and M, and, implicitly, the entire cell cycle duration^12^ (**Figure 1a**). Images were sampled every 10 minutes, and phase transitions were determined by eye. As expected, we found that G1 showed the greatest degree of variability across different cell lines, ranging from 2.1 hours in H9 to 7.9 hours in RPE. In contrast, the durations of S (7.6 h-10.1 h), G2 (3.4 h-4.0 h), and M (∼0.5 h) were relatively consistent across the three cell lines (**Figure 1b**). When looking among individual cells, however, both G1 and G2 showed substantial variability within each cell type (**Figure 1c-d**). S phase duration was the most tightly distributed phase^30^ with consistent coefficient of variance across different cell lines, even in the near-triploid U2OS. Thus, G1 is the most variable across different cell lines, but G1 and G2 are both highly variable within each cell line.

**Figure 1.**
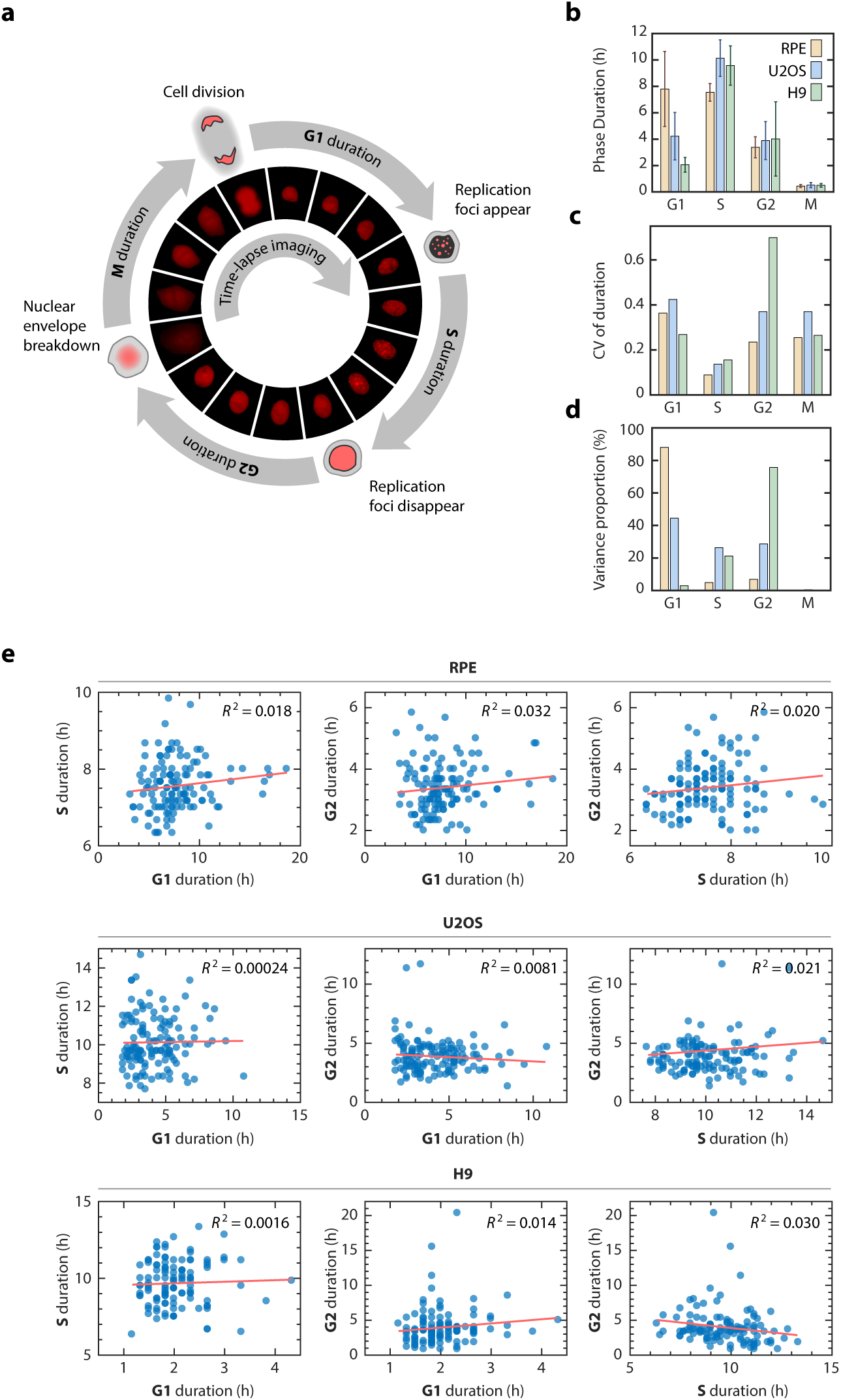
Variation and correlation among cell cycle phase durations in single cells. **a,** Diagram of a the cell cycle composed of G1, S, G2, and M phases. Phase durations were quantified by time-lapse fluorescence microscopy using a PCNA-mCherry reporter to identify 4 discrete events in the lifetime of an individual cell. Images were acquired every 10 min and transitions were identified and recorded by eye. The error of reported durations is ±5 min with standard deviation *σ* = 2.9 min. **b,** Mean phase durations in RPE, U2OS, and H9 cell lines. Error bars represent standard deviations. **c,** Coefficient of variation (CV) of phase durations. **d,** Percentage of the total variation in cell cycle duration contributed by individual phases. **e,** Correlations between individual cell cycle phase durations. Sample sizes were adequate to detect correlations (see Materials and Methods). *n* = 125 (RPE), 130 (U2OS), 113 (H9). *R^2^*, square of Pearson correlation coefficient.

We then asked whether any of the phase durations were correlated in individual cells. Correlation between phase durations would indicate the existence of “cellular memory” of the progression rate that persists for more than one cell cycle phase, as would be expected from previous studies^3,17^. We compared the durations of G1, S, and G2 phases only since the duration of M phase (∼30 mins) was significantly shorter than the other phases; similar to the image sampling rate (10 mins); and contributed little variance to the total cell cycle duration (**Figure 1d**)^17^. Surprisingly, we detected neither a significant (*P* < 0.01) nor strong (R^2^ > 0.1) correlation between any pair of phase durations under basal conditions (**Figure 1e, Figure S1**). This lack of correlation was not due to measurement error because we were able to readily detect correlations between phases in sister cells for every cell type (**Figure S2a**), as reported previously^2–4^. Furthermore, a power analysis revealed that our sample size would be adequate to detect significant correlations, if present (**Experimental Procedures**). We further note that many phases showed pronounced variability, indicating that the lack of correlation was not due to a lack of variability under basal conditions. Thus, contrary to previous claims that cell cycle phases are correlated, we find no evidence for phase coupling for these three human cell lines. This observation suggests that, under normal proliferating conditions, the effects of factors that determine the duration of each cell cycle phase are independent of that governing the duration of the previous, or next, cell cycle phase.

### Each cell cycle phase follows an Erlang distribution with a characteristic timescale and rate

The independence of phase durations suggested that each phase may be subject to a unique rate-governing process. We therefore examined the probability distributions of the phase durations in order to define the underlying stochastic processes driving them. All phases followed a similarly-shaped distribution characterized by a minimum duration time and skewed right tail (**Figure 2a**). This distribution immediately ruled out a one-step stochastic process, which would be expected to produce an exponential distribution of phase durations^1^. Instead, each distribution of phase durations resembled an Erlang distribution, which represents the sum of *k* Poisson processes with rate λ (**Figure 2b**). The Erlang distribution was originally developed to describe the waiting time before a series of telephone calls is handled by an operator^31^. In its application to the cell cycle, each phase can be thought of as a series of steps that proceeds at some fundamental rate^12^. The total amount of time needed to complete all steps in the phase has an Erlang distribution^32^. This model does not claim that each cell cycle phase is merely a series of precisely *k* steps. However, the Erlang model does provide a phenomenological description of cell cycle progression that has a simple and relevant biological interpretation: each cell cycle phase can be viewed as a multistep biochemical process that needs to be completed in order to advance to the next cell cycle phase. Similar models have been proposed to describe the “microstates” of stem-cell differentiation, a sequential biological process that undergoes similar discrete, observable state transitions at the same rate throughout the differentiation process^33^. In contrast to the differentiation process, model fitting suggested that a single rate parameter for all cell cycle phases was unable to fit the data well (**Figure S3a**), arguing that each cell cycle phase is controlled by distinct rate-governing mechanisms.

**Figure 2.**
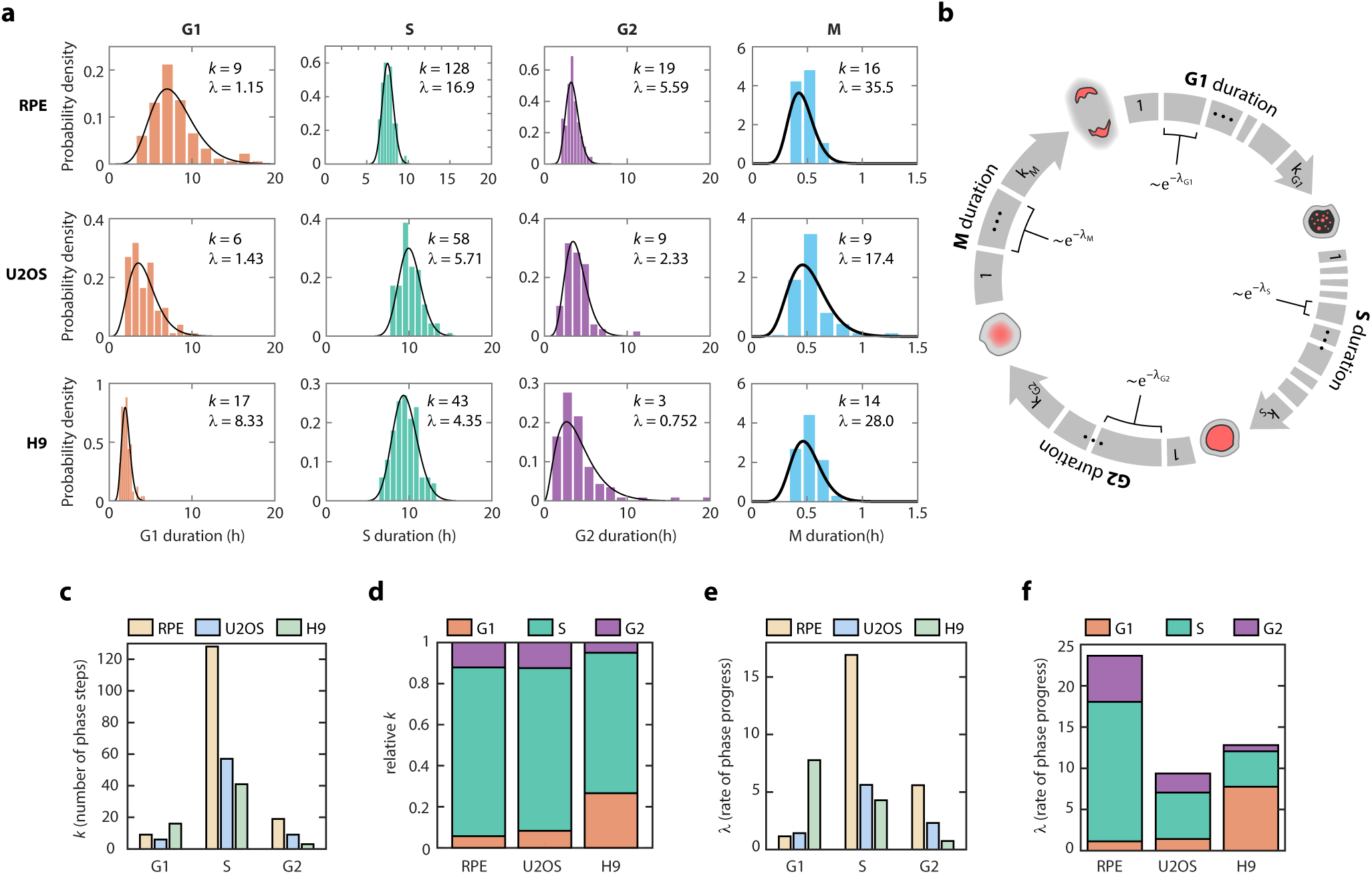
Erlang model of cell cycle progression. **a,** Distributions of cell cycle phase durations for RPE, U2OS, and H9 cells using single-cell measurements of phase duration reported in Figure 1. Black curves represent fits to Erlang distribution. **b,** Erlang model of cell cycle progression. Each phase consists of a distinct number of steps, *k*. Each step is a Poisson process with rate parameter, *λ*. After fitting each phase to the Erlang distribution, we were able to accurately simulate all phase durations except for M phase (2-sided Kilmogorov-Smirnov test for difference between measured and simulated distributions, **Figure S3b-c**). **c,** Fitted shape parameter, *k*, representing the number of steps for each phase. **d,** Normalized shape parameter, *k*, for G1, S, and G2 phases in RPE, U2OS, and H9 cells. Bar height represents the fraction of total cell cycle steps spent in each phase. **e**, Fitted rate parameter, *λ*, representing the progression rate of each step within a cell cycle phase. **f**, Rate parameter *λ* for each phase, shown by cell type.

By fitting the experimentally measured distributions of phase durations, we obtained two parameters for G1, S, G2, and M for each cell line: a shape parameter, *k*, which represents the number of steps within a phase; and a rate, λ, which represents how quickly on average the step is completed (**Figure 2a**, black curves). Using the estimated parameters, we were able to accurately simulate the cell cycle phase durations under basal conditions for all phases except for M phase, for which the time resolution of measurement was low (10 mins) compared to the average duration (∼30 mins) (**Figure S3b-c**). When we compared the shapes and rates across cell lines, several interesting points emerged. First, the number of steps was high (*k* = 43-128) for S phase but low for both G1 and G2 (*k*<20) (**Figure 2c**). In addition, although the absolute number of steps differed across cell types, the proportions of step for each phase were highly conserved, especially for RPE and U2OS (**Figure 2d**). This conserved proportionality suggests that a similar number of biochemical processes are involved in the same phase in each cell type. Second, the rate parameters generally followed the trends of the step parameters across cell lines, with high λ corresponding to high *k* (**Figure 2c, 2e**). This trend suggests that, regardless of the cell cycle phase, each cell line had a different strategy for cell cycle progression—RPE preferred to progress at higher rates through more numerous steps, followed by U2OS, then by H9 with slower rate and fewer step numbers. The one exception to this pattern was G1 in H9 (**Figure 2d, 2f**), which is consistent with the unusually short G1 duration in embryonic stem cells^7,34,35^. More generally, our analysis suggests that each cell cycle phase operates under independent control with respect to its rate of progress. This implies a memoryless property of cell cycle phases in which the rate of progress through a given phase does not depend on the rate of any previous phase.

### Cell cycle phase durations remain uncoupled when phase durations are altered

To further test whether phase durations are independently controlled, we introduced a series of perturbations aimed at altering the durations of specific phases and asked whether subsequent phases remained uncoupled. We introduced perturbations in the non-transformed RPE cell line, whose cell cycle regulation was unperturbed and was sufficiently resistant to perturbations^36^. We first specifically perturbed G1 length by inducing oncogene activation (**Figure 3a**). Overexpressing the oncogene Myc strongly and specifically shortened G1 by 55% (**Figure 3b, Figure S4a**) without introducing coupling among cell cycle phases on a single-cell level (**Figure 3c, Figure S4b**). We next targeted S phase by introducing replication stress with aphidicolin, and inhibitor of DNA polymerase (**Figure 3d**). Aphidicolin specifically prolonged S phase while leaving G2 duration unchanged (**Figure 3e, Figure S4c**), and there was no evidence of coupling between S phase and G2 (**Figure 3f, Figure S4d**). We next asked whether phases could become coupled by perturbing multiple phases. We prolonged all phases by incubating cells at 34°C (**Figure 3g**). Each phase lengthened by a similar proportion (**Figure 3h**, **Figure S4e**) instead of an absolute magnitude (**Figure S4f**). Surprisingly, even though all phases lengthened proportionally in response to lower temperature, the phase durations remained uncoupled at the single-cell level (**Figure 3i, Figure S4g**).

**Figure 3.**
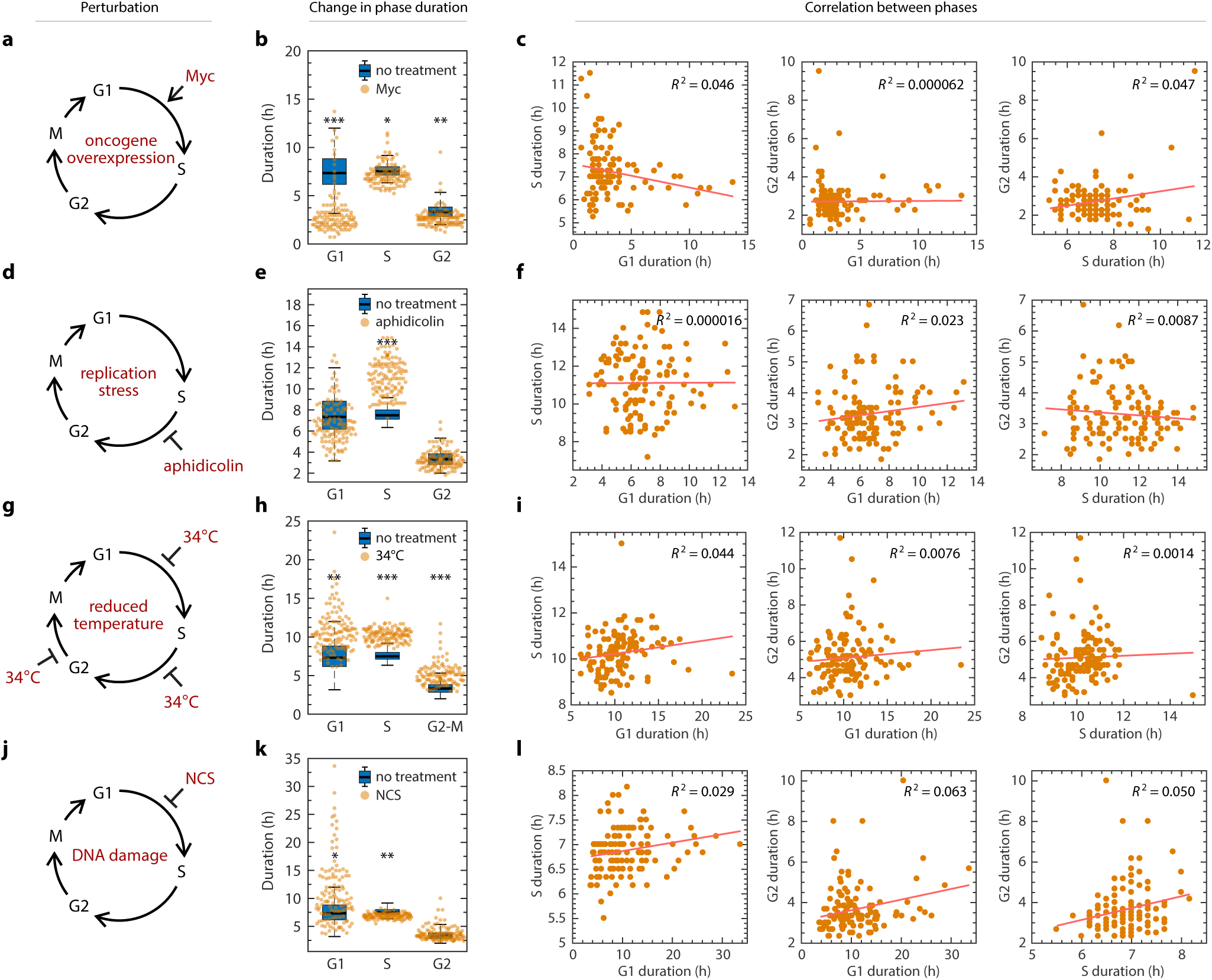
Lack of coupling among cell cycle phases under perturbation. **a,** Schematic of shortening G1 by Myc overexpression. RPE cells infected with retrovius harboring a taxmoifen-inducible Myc-overexpression construct. **b,** Shift in phase durations of RPE cells overexpressing Myc. **c,** Pairwise correlation between cell cycle phase durations of RPE cells overexpressing Myc. **d,** Schematic of prolonging S phase by replication stress using aphidicolin. Asynchronously proliferating RPE cells were treated with 50 ng/mL aphidicolin for 8 hours, washed with PBS, and then replenished with fresh media. Only cells whose S phase overlaped with the 8-hour treatment window were analyzed. **e,** Shift in phase durations of RPE cells treated with 50 ng/mL aphidicolin. **f,** Pairwise correlation between cell cycle phase durations under aphidicolin treatment. **g,** Schematic of prolonging all phases by incubating cells at 34°C. **h,** Shift in phase durations of RPE cells incubated at 34°C. **i,** Pairwise correlation between phase durations for cells incubated at 34°C. **j,** Schematic of prolonging G1 by DNA damage using NCS. Asynchronously proliferating RPE mother cells were treated with 25 ng/mL NCS, and their daughter cells were analyzed for a full cell cycle. **k,** Shift in phase durations of RPE cells treated with NCS. In panels **b**, **e**, **h**, and **k**, boxplots representing the distributions of phase durations in untreated cells are underlaid for comparison. *, *P* < 1 × 10^−5^; **, *P* < 1 × 10^−10^; ***, *P* < 1 × 10^−20^, 2-sided Kolmogorov–Smirnov test. Number of cells: Myc, *n* = 116; aphidicolin, *n* = 126; 34°C, *n* = 122; NCS, *n* = 119.

We next introduced the DNA damaging agent neocarzinostatin (NCS) to mother cells and measured the phase durations for daughter cells (**Figure 3j**). Recent work in human cells has shown that DNA damage signaling can persist through cell cycle phase transitions and lengthen the duration of G1, suggesting that coupling could potentially arise under genotoxic stress^11,23,37^. As expected, we found that DNA damage significantly lengthened G1 in the daughter cells (**Figure 3k, Figure S4h**). However, we found no strong correlations between phase durations, with the possible exception of G1 and G2, which showed a weakly significant correlation (R^2^ = 0.063, p = 0.006) (**Figure 3l, Figure S4i**). We then asked whether perturbing the duration of G2 in the mother could lead to a correlated G1 duration in the daughter cells. We found that DNA damage induced at different phases led to different responses in the daughter cells^12^ (**Figure S5a**) and that DNA damage in the mother cell’s S phase prolonged both the mother’s G2 phase and their daughters’ G1 phases in a dose-dependent manner (**Figure S5b**). However, these prolonged phase durations were still uncorrelated (**Figure S5c**), implying that the variations in phase duration were contributed by an external component (i.e. damage levels) and an intrinsic component, and that this intrinsic component did not couple phases. Therefore, whichever factors determined G2 duration did not necessarily determine G1 duration. Thus, although increasing levels of DNA damage increased both G2 and the subsequent G1, a mother with a very prolonged G2 did not necessarily have daughters with very long G1 durations, indicating that there was no phase coupling between G2 and the subsequent G1 on a single-cell level. We observed no effect of maternal G2 prolongation on the S and G2 phases in the daughter cells (**Figure S5a**). Taken together, these results suggest that although daughter cells can retain memory of maternal stress^23^, this memory is erased after committing to proliferation.

Our results suggest that the rates of progression for each cell cycle phase are controlled in a manner that leads to uncoupling between phase durations. We find no evidence of proportionality between phases as proposed by the stretched cell cycle model^3^, although we can reproduce the presence of a strong linear correlation between individual phase durations and the total cell cycle duration (**Figure S2b-c**). Further, we show that the Pearson correlation coefficient merely represents the proportion of variance explained by that variable, which was determined by the relative phase duration and its variance (**Figure 1d, Figure S2d**), and is therefore not an indication of coupling^17^, or proportional stretching between phases^3^. Rather, the variability in cell cycle duration depends simply on the relative length of each cell cycle phase. Interestingly, we found that, for the three cell lines that we studied, the influence of combined S-G2-M ranged from accounting for a small part of total variability in RPE, to accounting for the majority of total variability in H9 (**Figure S2e**). Thus, the RPE cell line is more consistent with the Smith-Martin model in which G1 accounts for most of the variability in cell cycle length^1^. In contrast, rapidly proliferating H9 cells are most similar to the lymphocytes that form the basis for the stretched model^3^, in which variability in cell cycle duration stemmed primarily from S and G2.

### A model for heritable factors governing independence of phase duration

We next sought to reconcile our model with previous observations concerning the heritability of cell cycle phase durations. It has long been known that sibling cells are highly correlated in their total cell cycle durations as well as in the durations of individual phases^2–4,9^. These observations strongly suggest that there are heritable molecular factors that influence the rate of cell cycle progression. But this observation raises an obvious paradox: if cells retain factors that control the durations of cell cycle phases, how can consecutive phases be uncoupled and memoryless? To reconcile these two observations, we considered three models in which heritable factors influence the phase progression rates, and implicitly the phase durations. In the first model, which we refer to as the “one-for-all” model, one heritable factor influences the duration of all phases (**Figure 4a**). Under this model, all phases are strongly correlated as each phase shares common control over rate progression. However, the uncoupling between phases we observed (**Figures 1e, Figure 3**) is inconsistent with this model.

**Figure 4.**
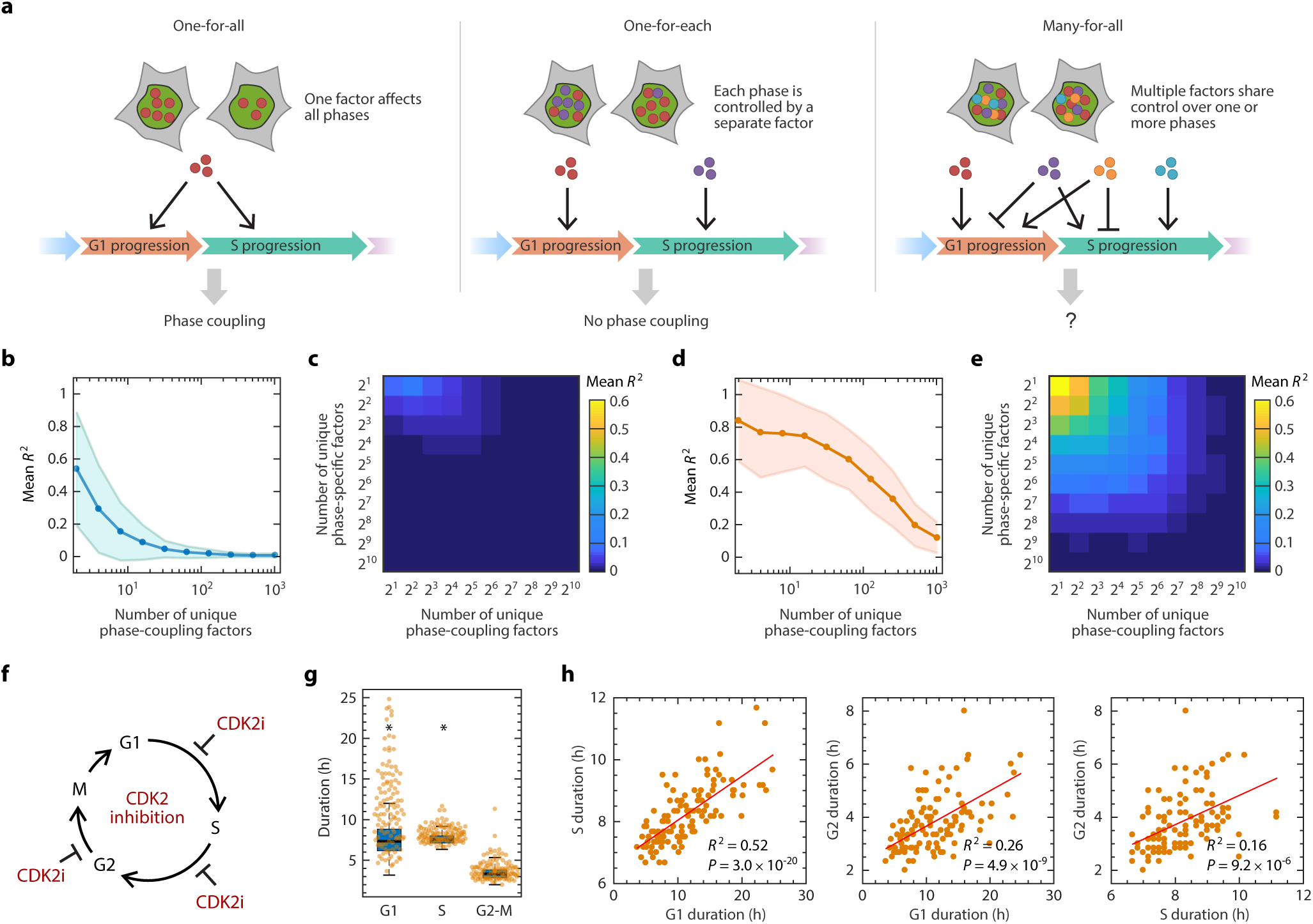
A model for heritable factors governing the rate of cell cycle phase progression. **a,** Alternative models for inheritance of molecular factors governing the durations of cell cycle phases. **b,** Simulation of the “strength of coupling” as a function of the number of unique phase-coupling molecule types under the many-for-all model. Each simulation generated 200 cells for which an *R* ^2^ value was calculated. *R* ^2^ values were averaged across 200 simulations. The shaded area represents the standard deviation of *R* ^2^ across the simulations. **c,** Simulation of coupling strength as a function of the number of unique phase-coupling and phase-specific factors. Phase-coupling factors have shared control over a pair of cell cycle phases, whereas phase-specific factors affect only one cell cycle phase. Strength of coupling is represented by mean *R* ^2^ value as in panel **b**. **d**, same as in **b**, but simulating the effect of perturbing a single phase-coupling factor by significantly increasing its abundance or activity. Perturbation was simulated by increasing the abundance of a phase-coupling factor by 10 fold. **e,** Same as in **c**, but simulating the effect of increasing a phase-coupling factor by 10 fold (see **Method Details**). **f,** Schematic of prolonging all phases by inhibition of CDK2. RPE cells were treated with 2 µM CVT-313 and the durations of each phase were quantified for a full cell cycle. **g**, Shift in phase durations for RPE cells treated with 2 µM CVT-313. A boxplot representing the distribution of durations in untreated cells is underlaid for comparison. *, *P* < 1 × 10^−5^, 2-sided Kolmogorov–Smirnov test. (*n* = 117 cells). **h**, Pairwise correlation between cell cycle phase durations upon treatment with CVT-313.

A second model, called “one-for-each”, entails that each cell cycle phase has its own rate-determining factor, and that these heritable factors propagate independently to daughter cells (**Figure 4a**). Under the one-for-each model, each cell cycle phase progresses under its own control independently of the previous phases, which is consistent with our results. However, this model contradicts well-established knowledge about several molecular factors that share control over multiple phase durations. For example, cells that have high E2F activity, which controls both the entry into S phase as well as DNA replication, are expected to progress through both G1 and S faster^19,20^. In contrast, cells with high Cdt1 expression, which functions to license origins for replication, finish G1 early but have a prolonged S phase^38,39^. Thus, we considered the possibility that there may be numerous types of heritable factors, each of which affects multiple phase durations in potentially different directions. We called this the “many-for-all” model (**Figure 4a**). Under this model, each phase is under shared control by multiple types of factor. Because each factor individually has a coupling effect, the net effect of a group of such factors could potentially lead to coupling of cell cycle phase durations.

To explore whether the heritable factors would lead to phase coupling under the many-for-all model, we computationally modeled the coupling between two phases under shared control as a function of numbers of unique factor types (**Method Details**). Simulation results revealed that the coupling between phases weakened as the number of unique coupling factor types increased (**Figure 4b**). Intuitively, this uncoupling effect arises as the net effect of numerous heritable factors dilutes the effect of individual coupling factors, preventing any single coupling factor from dominating control over phase durations. In addition, introducing more phase-specific factors, which only affect a single phase, would further uncouple the phases by diluting the coupling factors’ effects (**Figure 4c**). Because we observed no correlation between cell cycle phase durations under basal or perturbed conditions, our experimental results are consistent with the regime of numerous factor types under many-for-all model of cell cycle phase progression.

We gained further insight into the inheritance of phase-coupling factors by analyzing sister cell pairs. Because sister cells share similar amounts of heritable factors due to shared cytoplasmic and genetic content^40,41^, all of the models above would be expected to produce correlations between sister cells’ phase durations. However, in order to achieve the observed phase uncoupling in individual cells, the distribution of each type of heritable factor to daughter cells must be independent of the others (see **Method Details**). If factors segregate independently, then even in sister cells—for which phase durations are highly correlated— the noise for each cell cycle phase length is expected to be uncoupled between sisters. For example, the differences between G1 durations in sisters would not be expected to correlate with differences between S durations. To support this hypothesis, we show that even though cell cycle phase durations are highly correlated between sisters (**Figure S2a**), we found no correlation between the differences in sibling cells’ phase durations for any pair of phases (**Figure S6a**). Thus, phase durations appear to be controlled by a large number of heritable factors that segregate independently during cell division.

### Perturbation of a single factor leads to coupling between cell cycle phase durations

According to the many-for-all model of heritable factors, no single factor dominates the coupling effect among phase durations. Therefore, we hypothesized that cell cycle phases could be coupled by increasing the levels, or activity, of a molecular factor that controls more than one phase so that the effect of this factor becomes dominant. To explore this possibility, we computationally simulated perturbing a single factor by significantly increasing its abundance (**Method Details**). Consistent with our hypothesis, increasing one factor’s net effect introduced coupling between phase durations (**Figure 4d-e**). To experimentally test this prediction, we introduced the cyclin-dependent kinase 2 (CDK2) inhibitor CVT-313^42^ and measured the resulting phase durations (**Figure 4f**). CDK2 is a key cell cycle regulator involved in multiple cell cycle phases, including G1/S transition, S phase progression, and G2/M transition^43–46^. Adding the CDK2 inhibitor mimics the effect of increasing the levels of p21, a potent negative regulator of CDK2. As expected, CDK2 inhibitor treatment prolonged all cell cycle phases, having the strongest effect on G1 (**Figure 4g**, **Figure S6b**). Moreover, consistent with the model’s prediction, inhibiting CDK2 also introduced a strong correlation between each pair of phase durations (**Figure 4h, Figure S6c**). It is notable that we were able to force correlations by inhibiting CDK2, a single molecular factor that affects multiple phases, but not by decreasing temperature, which—although also influencing all phases—perturbed a large number of molecular factors’ activity.

### Uncoupling between cell cycle phases is disrupted in a cancer cell line

The above results in a non-transformed cell line suggest that cell cycle phases are not intrinsically uncoupled, but only appear uncoupled due to a large number of factors sharing control over multiple phases. It is possible that this balance of molecular control is disrupted in other cell lines bearing either excessive or depleted abundance of cell cycle control factors. To examine phase coupling in cells with dysregulated cell cycle control, we measured whether coupling occurred after perturbing specific phases in U2OS cells, which are known to have a perturbed G1 checkpoint^26,27^. We used NCS to induce DNA damage in mother cells and quantified the daughter cells’ phase durations (**Figure 5a**). As in RPE cells, only G1 was significantly prolonged by the DNA damage induced in the mother cells (**Figure 5b, Figure S7a**). However, unlike in RPE cells, DNA damage induced in U2OS mother cells introduced coupling between daughters’ G1 and S durations and resulted in a positive correlation (**Figure 5c, Figure S7b**), even in the absence of a increase in S phase duration at the population level. We next perturbed S phase with aphidicolin to induce replication stress in U2OS cells (**Figure 5d**). As in RPE cells, we observed significant prolongation of S phase duration only (**Figure 5e, Figure S7c**). In contrast to RPE cells, however, S phase lengthening was coupled to longer durations of both G1 and G2 (**Figure 5f, Figure S7d**). Interestingly, the duration of G1—which elapsed before treatment of cells in S phase and was therefore unaffected—predetermined a cell’s sensitivity to aphidicolin: longer G1 duration correlated with a longer perturbed S phase. This result suggested that one or more phase-controlling factors contribute both to G1 and S phase progression but only become rate-limiting for S phase progression under replication stress. In contrast, the many-for-all model predicted that a perturbation promiscuously affecting many factors would not introduce coupling despite a dysregulated cell cycle. Consistent with this prediction, low temperature perturbation prolonged cell cycle phases without introducing phase coupling (**Figure S7e-i**). Taken together, these results suggested that in U2OS cells, coupling between cell cycle phases became apparent under certain stressed conditions.

**Figure 5.**
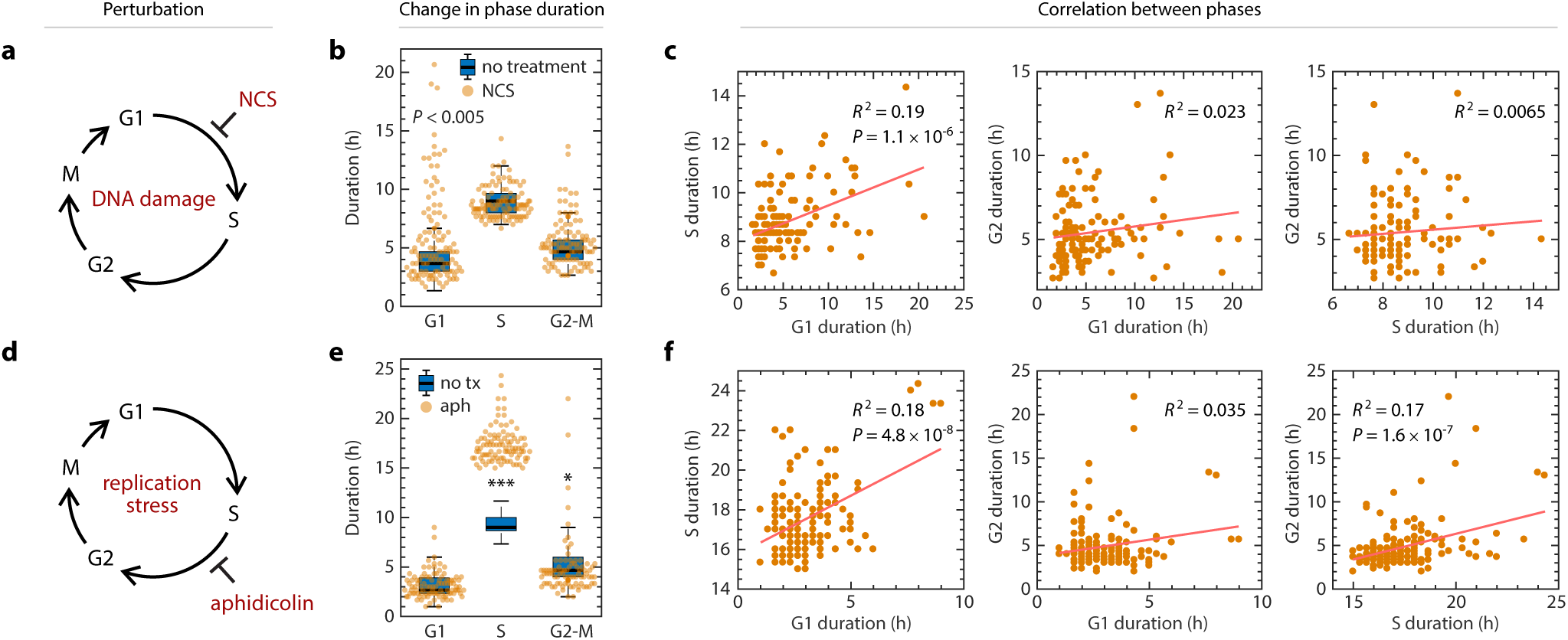
Stress-induced coupling of cell cycle phases in a cancer cell line. **a,** Schematic of prolonging G1 by DNA damage using NCS. Asynchrnously proliferating U2OS mother cells were treated with 100 ng/mL NCS, and their daughter cells were analyzed for a full cell cycle. **b,** Shift in phase durations of U2OS cells treated with NCS. **c,** Pairwise correlation between phase durations for U2OS cells treated with NCS. **d,** Schematic of prolonging S phase by replication stress using aphidicolin. Asynchronously proliferating U2OS cells were treated with 50 ng/mL aphidicolin for 8 hours, washed with PBS, and then replenished with fresh media. Only cells whose S phase overlaped with the 8-hour treatment window were analyzed. **e,** Shift in phase durations of U2OS cells treated with 50 ng/mL aphidicolin. **f,** Pairwise correlation between cell cycle phase durations under aphidicolin treatment. In panels **b** and **e**, boxplots representing the distributions of phase durations in untreated cells are underlaid for comparison. *, *P* < 1 × 10^−5^; ***, *P* < 1 × 10^−20^, 2-sided Kolmogorov–Smirnov test. Number of cells: NCS, *n* = 114; aphidicolin, *n* = 153.

## Discussion

The ability to monitor single cells in real time has revealed the integral role of the cell cycle in controlling multiple cellular processes including the decision between proliferation or quiescence^19,47,48^, differentiation of pluripotent cells into multiple lineages^49–51^, and susceptibility to chemotherapeutics^52^. Therefore, knowledge of the principles underlying cell cycle progression will provide insights into how cell fate decisions are determined and how they can be manipulated for therapeutic gain. In this study, we developed a phenomenological model for cell cycle progression using precise measurements for G1, S, G2, and M durations in three human cell lines. We found that cells with functional cell cycle checkpoints progress through each phase at an independent rate that is independent of previous phases. This independence between phases, which can be modeled as a sequence of memoryless processes, can be explained by the presence of many molecular factors that each contributes a small effect over one or more phase durations.

The lack of correlation between phase durations was unexpected and would seem to disagree with previous experimental results and theoretical models. We offer several explanations for this discrepancy. First, many previous data supporting cell cycle phase coupling relied on correlations between the total cell cycle duration and a part of the total duration^3,17^. Such a relationship does not necessarily imply coupling (**Figure S2b-e**). In contrast, we directly measured the degree of coupling between individual phases and found no evidence of significant coupling. Second, previous studies examined different cell types (e.g., mouse lymphocytes) that could have different profiles of factors controlling cell cycle progression such as elevated endogenous DNA damage levels^53^ or different roles of tumor suppressors p53 and Rb in permanent arrest^54^. According to the many-for-all model, observation of coupling between cell cycle phases would imply that phase-controlling factors are relatively more abundant in certain cell types. Third, we employ a more accurate method of measuring cell cycle phase durations based on appearance and disappearance of PCNA foci during S phase^12^. Previous studies employed the FUCCI reporter system to distinguish G1 and S-G2-M cells, but this system is known to give unclear cell cycle phase boundaries^55^. Fourth, it is possible that previous studies analyzed a mixed population of cells (e.g., cells at different stages of differentiation or maturation) that could result in a correlation between phase durations across the different cell types^56^. To avoid this problem, we analyzed three clonal cell lines under steady state growth conditions. Finally, certain stressful growing conditions may introduce coupling between cell cycle phases, such as high levels of environmental stress (**Figure 5c, 5f**) or upregulation of potent cell cycle inhibitors such as p21, which inhibits CDK2 activity (**Figure 4h**)^57^.

Our model explains why phase coupling may not be observed despite the fact that coupling factors are known to exist. For example, a sizer model for cell cycle control, in which the cell size controls cell cycle progression by G1/S and G2/M transitions, would predict correlation between durations of cell cycle phases in single cells because cell size is inherited within the same cell cycle^58–60^. Similarly, a CDK activity threshold mode, in which cell cycle phase progression is achieved by ordered substrate sensitivity, would also predict coupling between cell cycle phases since CDK activity is an intrinsic factor within the same cell that acts throughout the cell cycle^61,62^. However, according to the many-for-all model, in which each phase-controlling factor has shared control over multiple phases, the net effect of many such factors leads to uncoupling between cell cycle phase durations due to large number effect. Under this model, perturbations that influence more phase-length factor types are expected to show less coupling, whereas perturbations that influence one or a few factors are expected to introduce stronger coupling. Consistent with this prediction, we find the degree of coupling is the weakest under the most phase-specific perturbation with replication stress (**Figure 3f**) and under the most promiscuous perturbation of reduced temperature (**Figure 3i**), in which presumably all biochemical processes are slowed. Myc overexpression leads to global transcription changes involving broad spectrum of cellular functions^63,64^, and DNA damage induces the DNA damage response (DDR) network, which is an ensemble of components^65^. Both perturbations involve large number of phase-controlling factors that can preserve the diversity of factors and lead to mild phase coupling. In contrast, perturbing a single phase-controlling factor by CDK2 inhibition introduces the strongest coupling among all cell cycle phases. This model is consistent with the observation that induced lengthening of one gap phase in *Drosophila* leads to accelerated progress through the subsequent gap phase via E2F1 regulation^22^, although further work is required to determine whether the phases are actually coupled in single cells. Thus, our model harmonizes with other descriptions of cell cycle progression by providing a framework for predicting when phase couplings occur.

Phase coupling may also be an indicator of dysregulated cell cycle control. We found that in the U2OS cell line, which harbors known defects in its G1 checkpoint^26,27,66^, stresses such as DNA damage and replication stress introduced phase coupling. Stress signals such as ATM and ATR are phase-controlling factors that negatively regulate the cell cycle progression. Functional checkpoints normally detect stress signals and wait for the signals to resolve before allowing cell cycle progression to resume, making cell cycle phases effectively “insulated” from the previous phase, which could contribute to the memoryless cell cycle phases. Without a fully functional checkpoint, the memory of stress signal levels could be transmitted from one phase to the next and lead to phase-coupling^67,68^. The coupling may also be explained by oncogenic changes in cancerous cells, which are often accompanied by oncogene activation and tumor suppressor loss. Oncogene activation leads to overexpression and thus dominance of a few phase-controlling factors, whereas tumor suppressor loss decreases the pool of phase-controlling factor types^69^, both of which could lead to imbalance in the competing pool of factors and susceptibility to phase coupling under stress. Cancer cells are characterized by genome instability and defective DDR pathways. Therefore, cancer cells are over-reliant on the remaining intact part of the DDR network such as ATM and Chk1^70–72^. When further DNA damage or replication stress is incurred, these few critical components are further induced, which may lead to dominating control over other factors and thus causing phase-coupling. Further work is required to determine whether phase coupling is a common feature of cancer cell lines.

Finally, our work is consistent with the observation that the memory of cell cycle duration is lost when a cell divides, as evidenced by the lack of correlation between mother and daughter cells’ cell cycle durations^4,24,73^. Sandler *et. al* show that this apparent stochasticity is in fact controlled by underlying deterministic factors that operate on a time scale different than the cell cycle control. They propose a “kicked” model in which an out-of-phase external deterministic factor leads to lack of correlation between consecutive cell cycles. Consistent with these observations, our results suggest that, in cells with intact cell cycle regulations, memory of cell cycle phase durations is not only lost over generations but also within a single cell’s lifetime between consecutive cell cycle phases. Thus, the emerging theme for governance of cell cycle progression is that phases are strongly coupled between temporally concurrent cells (i.e., sisters and cousins) but not coupled longitudinally over time.

## Experimental Procedures

### Cell Culture

hTERT retinal pigment epithelial cells (RPE) were obtained from the ATCC (ATCC^®^ CRL-4000^™^) and cultured in DMEM medium supplemented with 10% fetal calf serum (FBS) and penicillin/streptomycin. U2OS cells were obtained from the laboratory of Dr. Yue Xiong and cultured in DMEM medium supplemented with 10% fetal bovine serum (FBS) and penicillin/streptomycin (Gibco). WA09 (H9) hES cell line was purchased from WiCell (Wisconsin) and maintained in mTeSR1 (85850, StemCell Technologies) on growth factor reduced Matrigel (354230, BD). Cells were passaged using Trypsin (25300054, Gibco) for RPE and U2OS or ReLeSR™ (05872, StemCcell Technologies) for H9 as needed. When required, the medium was supplemented with selective antibiotics (2 μg/mL puromycin for RPE and U2OS; 0.5 μg/mL puromycin for H9) (A1113803, Gibco).

### Chemical and Genetic Perturbation of the Cell Cycle Phases

For NCS treatment, medium was replaced with fresh medium supplemented with neocarzinostatin (N9162, Sigma-Aldrich) during experiments. For myc overexpression, RPE cells were infected with fresh retrovirus containing MSCV-Myc-ER-IRES-GFP and 1 μL polybrene. Cells were subsequently passaged post 48 hours of infection and seeded onto a glass-bottom plate for imaging. 16 hours prior to imaging, tamoxifen was added at a final concentration of 50 nM. For aphidicolin treatment, medium was replaced with fresh medium supplemented with aphidicolin (A0781, Sigma-Aldrich) for 8 hours during experiments, washed off once with PBS, and then replenished with imaging media described below. For CDK2 inhibition, cells were treated with 2 μM CVT-313 (221445, Santa Cruz) prior to starting the imaging.

### Cell Line Construction

The construction of the pLenti-PGK-Puro-TK-NLS-mCherry-PCNA plasmid was described in our previous publication^12^. The plasmid was stably expressed into RPE, U2OS, and H9 cells by first transfecting the plasmid into 293T cells to generate replication-defective viral particles using standard protocols (TR-1003 EMD Millipore), which were used to stably infect the RPE, U2OS, and H9 cell lines. The cells were maintained in selective media and hand-picked to generate a clonal population.

The MSCV-Myc-ER-IRES-GFP was made by cloning the Myc-ER from pBabe-puro-Myc-Er into MSCV-IRES GFP. pBabe-puro-myc-ER was a gift from Wafik El-Deiry (Addgene plasmid # 19128)^74^. MSCV-IRES-GFP was a gift from Tannishtha Reya (Addgene plasmid # 20672). The cloned plasmid was then sequenced and verified.

### Time-Lapse Microscopy

Prior to microscopy, RPE and U2OS cells were plated in poly-D-lysine coated glass-bottom plates (Cellvis) with FluoroBrite™ DMEM (Invitrogen) supplemented with 10% FBS, 4 mM L-glutamine, and penicillin/streptomycin. H9 cells were plated in Matrigel coated glass-bottom plates with phenol red-free DMEM/F-12 (Invitrogen) supplemented with 1x mTeSR1 supplement (85852, StemCell Technologies). Fluorescence images were acquired using a Nikon Ti Eclipse inverted microscope with a Nikon Plan Apochromat Lambda 40X objective with a numerical aperture of 0.95 using an Andor Zyla 4.2 sCMOS detector. In addition, we employed the Nikon Perfect Focus System (PFS) in order to maintain focus of live cells throughout the entire acquisition period. The microscope was surrounded by a custom enclosure (Okolabs) in order to maintain constant temperature (37°C) and atmosphere (5% CO2). The filter set used for mCherry was: 560/40 nm; 585 nm; 630/75 nm (excitation; beam splitter; emission filter) (Chroma). Images were acquired every 10 minutes for RPE and H9 cells and every 10 or 20 minutes for U2OS cells in the mCherry channel. We acquired 2-by-2 stitched large image for RPE cell. NIS-Elements AR software was used for image acquisition and analysis.

### Image Analysis

Images were sampled every 10 minutes. Image analysis on the cell cycle phase was performed by manually tracking each cell and recording the frame at which PCNA foci appeared (G1/S) or disappeared (S/G2) and nuclear envelope breakdown (G2/M) using ImageJ to quantify the durations of each cell cycle phase. This provided reliable measurement of phase durations with a measurement error of one time frame (±10 min). In addition, due to the nature of time-lapse imaging, there was an uncertainty regarding when the phase transition occurred within the 10-min time frame.

### In silico Mapping of Cell Cycle Progression in Individual Cells

We quantified the cell cycle phase durations of our cell lines by imaging asynchronously dividing cells. During the entire life of each individual cell, we took five time point measurements: the time of cell birth (tbirth), the onset of S phase (ts_onset), the end of S phase (ts_end), the time of nuclear envelope breakdown (NEB, tm_start), and the time of telophase (ttelophase), which were manually identified from the PCNA-mCherry reporter. These five time points allowed for quantifying the durations of four cell cycle phases: G1, S, G2, and M phases.

### Statistical Analysis and Sample Size

Sample size was calculated based on Type I error rate of 0.2, Type II error rate of 0.01, and R^2^ =0.1 to prevent false negative correlation, which resulted in 112 cells per condition^75^. Non-parametric bootstrap was performed with 10,000 iterations to calculate the distribution of correlation coefficients for each condition and the percentage of iterations with no significant correlation (R^2^ < 0.1).

## Acknowledgements

We thank Samuel Wolff for guidance on experiments and microscopy; Katarzyna Kedziora both for valuable discussions and sharing unpublished results that confirm the lack of phase correlation; Rebecca Ward for critical feedback on the manuscript; Gavin Grant for providing scientific insights on CDK2 inhibition; Po-Hao Huang for brainstorming ideas; and members of the Purvis Lab for helpful discussions and technical suggestions. This work was supported by the National Institutes of Health research grant DP2-HD091800 (J.E.P.) and training fellowship F30-CA213876 (H.X.C.); a Medical Research Grant from the W.M. Keck Foundation (J.E.P.); the Loken Stem Cell Fund; and the North Carolina University Cancer Research Fund.

## Author Contributions

H.X.C. constructed the PCNA-mCherry reporter cell lines. H.X.C. and J.E.P. designed the experiments. H.X.C., R.I.F., H.K.S., and R.J.K. performed live-cell imaging and experiments. H.X.C., R.I.F., H.K.S., and R.J.K. conducted image analysis and cell tracking. H.X.C. performed computational modeling and analysis. H.X.C. wrote the manuscript with contributions from all authors.

---

**Figure S1.**
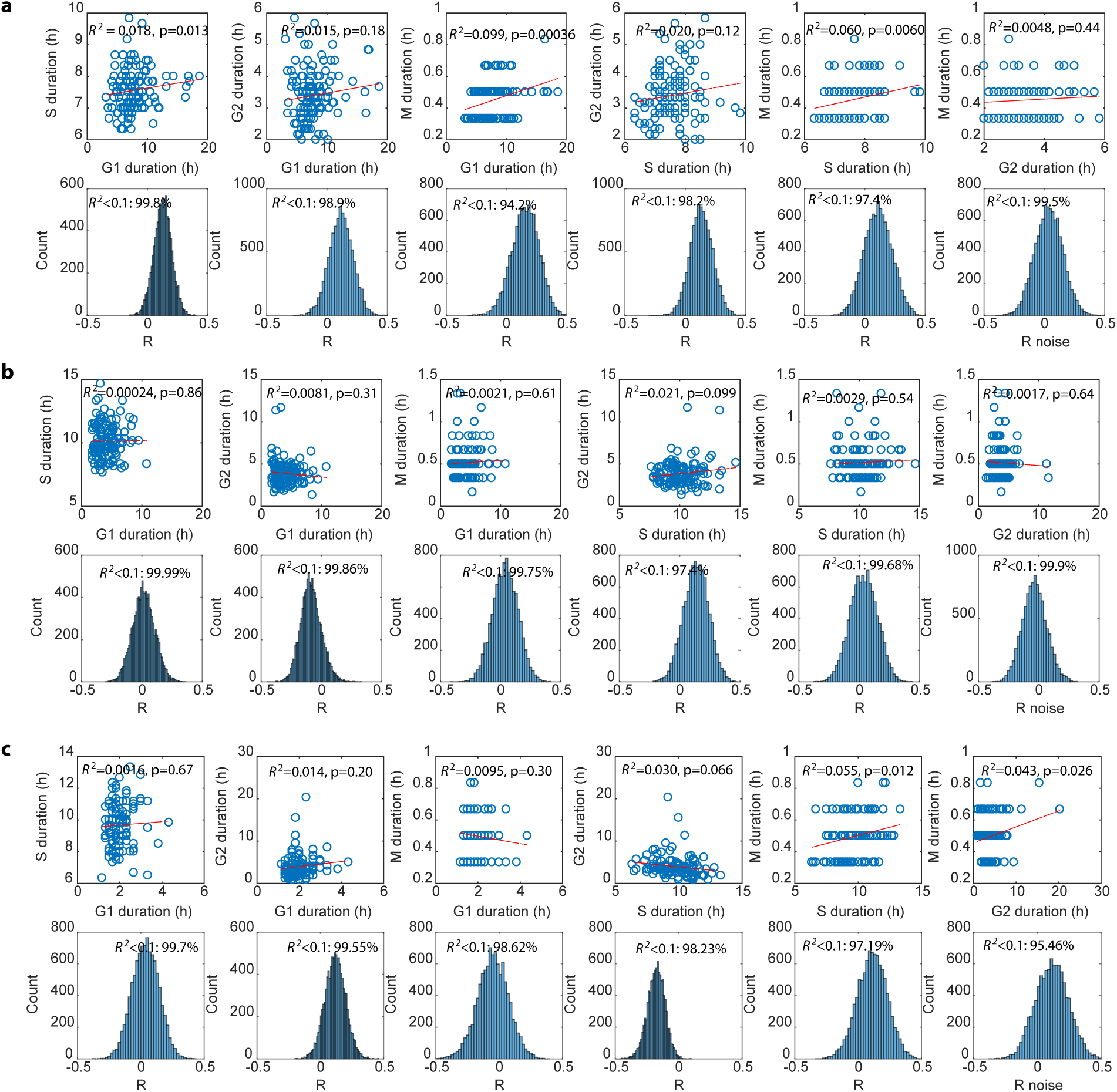
Pairwise correlations between cell cycle phases in three human cell lines. **a,** Upper panel: correlation between the cell cycle phase durations in RPE. Lower panel: Non-parametric bootstrap of the distribution of correlation coefficient (R), with consideration of measurement error (see Method details). n=10,000. **b,** Same as **Figure S5a**, but in U2OS. **c,** Same as **Figure S5a**, but in H9.

**Figure S2.**
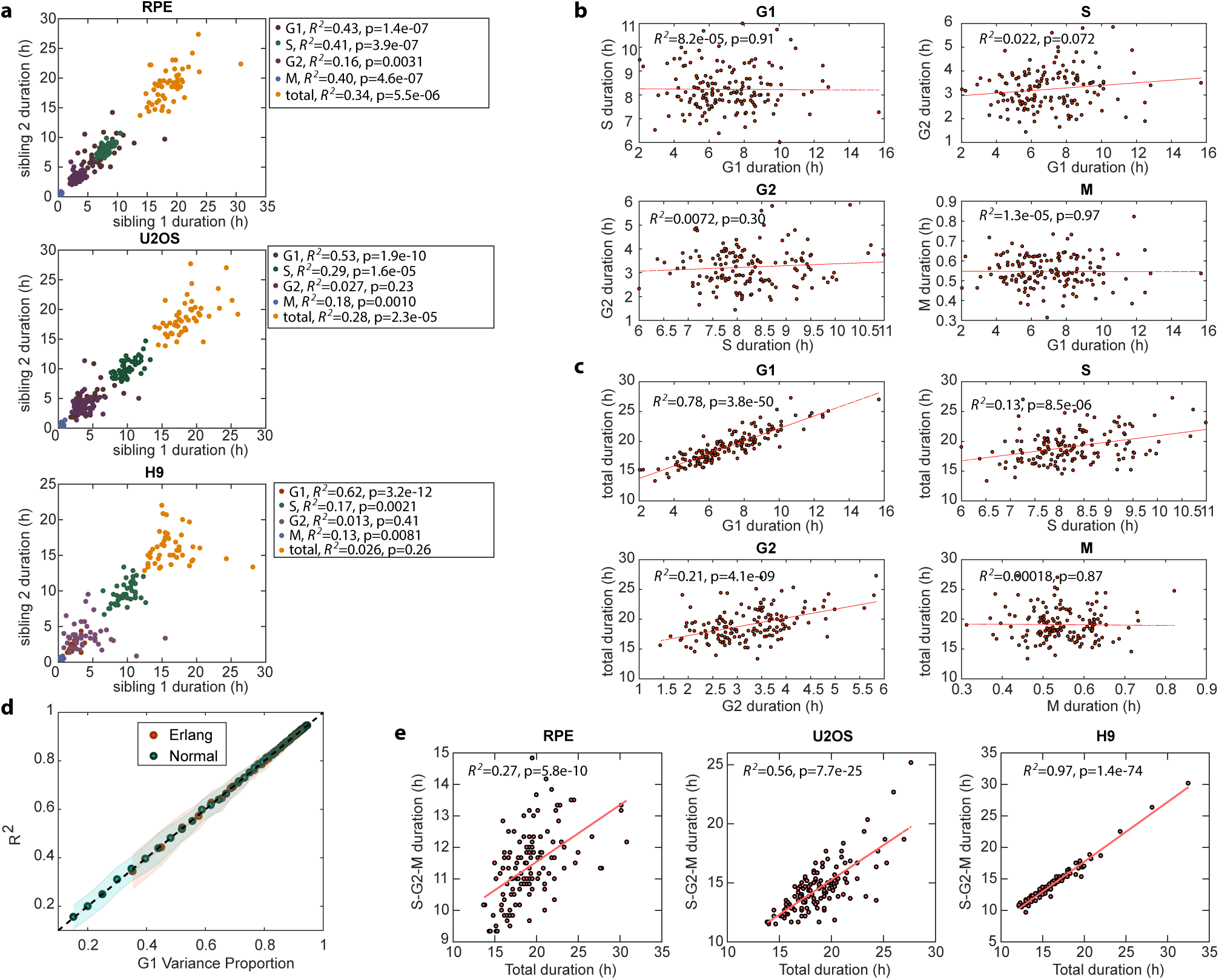
Correlation in cell cycle phase durations and its origin. **a,** Correlation between the cell cycle phase durations of sibling cells in RPE (upper panel), U2OS (middle pannel), and H9 (lower panel). Data were fit with linear regression, and Spearman correlation coefficients were calculated. n>100. **b,** Simulation of pairwise correlation between cell cycle phase durations under the Erlang model. The parameters were from the fittted RPE cell data in Figure 2a. **c,** Simulation of correlation between each cell cycle phase and the total cell cycle durations under the Erlang model, as in **Figure S2b**. **d,** Simulation of correlation coefficients as a function of the variance in total cell cycle duration contributed by G1. Data were simulated either under the Erlang model, as in **Figure S2b-c**, or under the normal distribution model. For the normal distribution model, parameters were choosen according to the mean and variance of the cell cycle phase duration’s distributions. The dashed line represents the diagonal line. **e,** Correlation between the combined S-G2-M phase and the total cell cycle duration in RPE (left), U2OS (middle), and H9 (right) cells’ experimental data. Data were fitted with linear regression.

**Figure S3.**
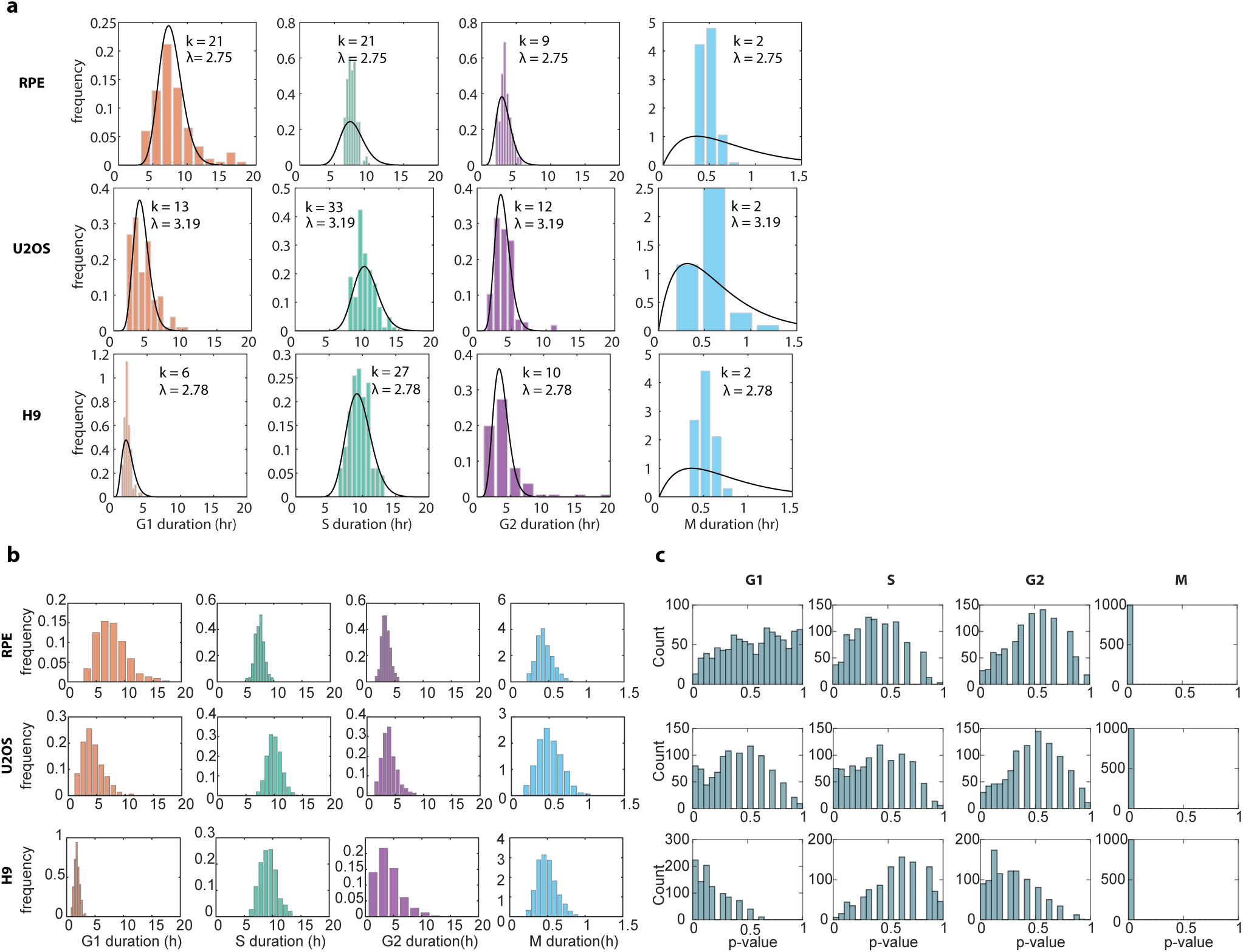
Fitting with a single rate parameter for all phases is insufficient to recapitulate the cell cycle distribution. **a,** Distributions of cell cycle phase durations fitted with a simple Markovian model with a single rate parameter. **b,** Simulations of the cell cycle phase durations of the Erlang model based on the fitted parameters. **c**, Distribution of the p-value, based on the Kolmogorov–Smirnov test, for significant difference in the cell cycle phase distributions between the experimental data and the simulated data from the Erlang model. For each simulation, 200 cells were generated, with a total of 1000 simulations.

**Figure S4.**
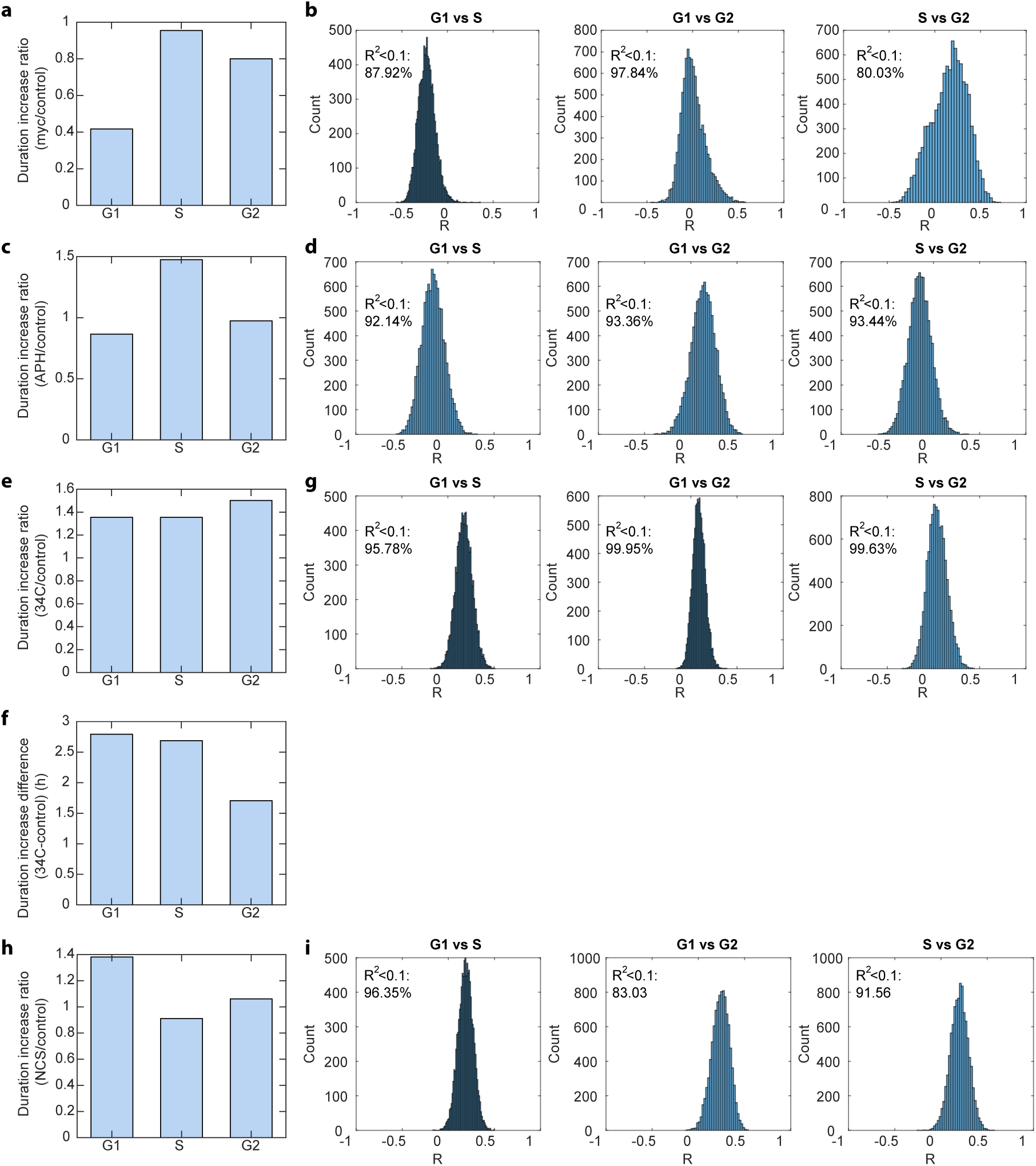
Perturbing cell cycle phase durations and phase coupling in RPE cells. a, Ratio of RPE’s cell cycle phase duration increase with myc overexpression relative to control. **b,** Non-parametric bootstrap of the distribution of correlation coefficient (R) for RPE under myc overexpression. n=10,000. **c,** Ratio of RPE’s cell cycle phase duration increase under 50 ng/mL APH treatment relative to control. **d,** Non-parametric bootstrap of the distribution of correlation coefficient (R) for RPE under 50 ng/mL APH. n=10,000. **e,** Ratio of RPE’s cell cycle phase duration increase at 34°C relative to control (37°C). **f,** Absolute magnitude of RPE’s cell cycle phase duration increase at 34°C relative to control (37°C). **g,** Non-parametric bootstrap of the distribution of correlation coefficient (R) for RPE at 34°C. n=10,000. **h,** Ratio of RPE’s cell cycle phase duration increase under 25 ng/mL NCS treatment relative to control. **i,** Non-parametric bootstrap of the distribution of correlation coefficient (R) for RPE under 25 ng/mL NCS treatment. n=10,000.

**Figure S5.**
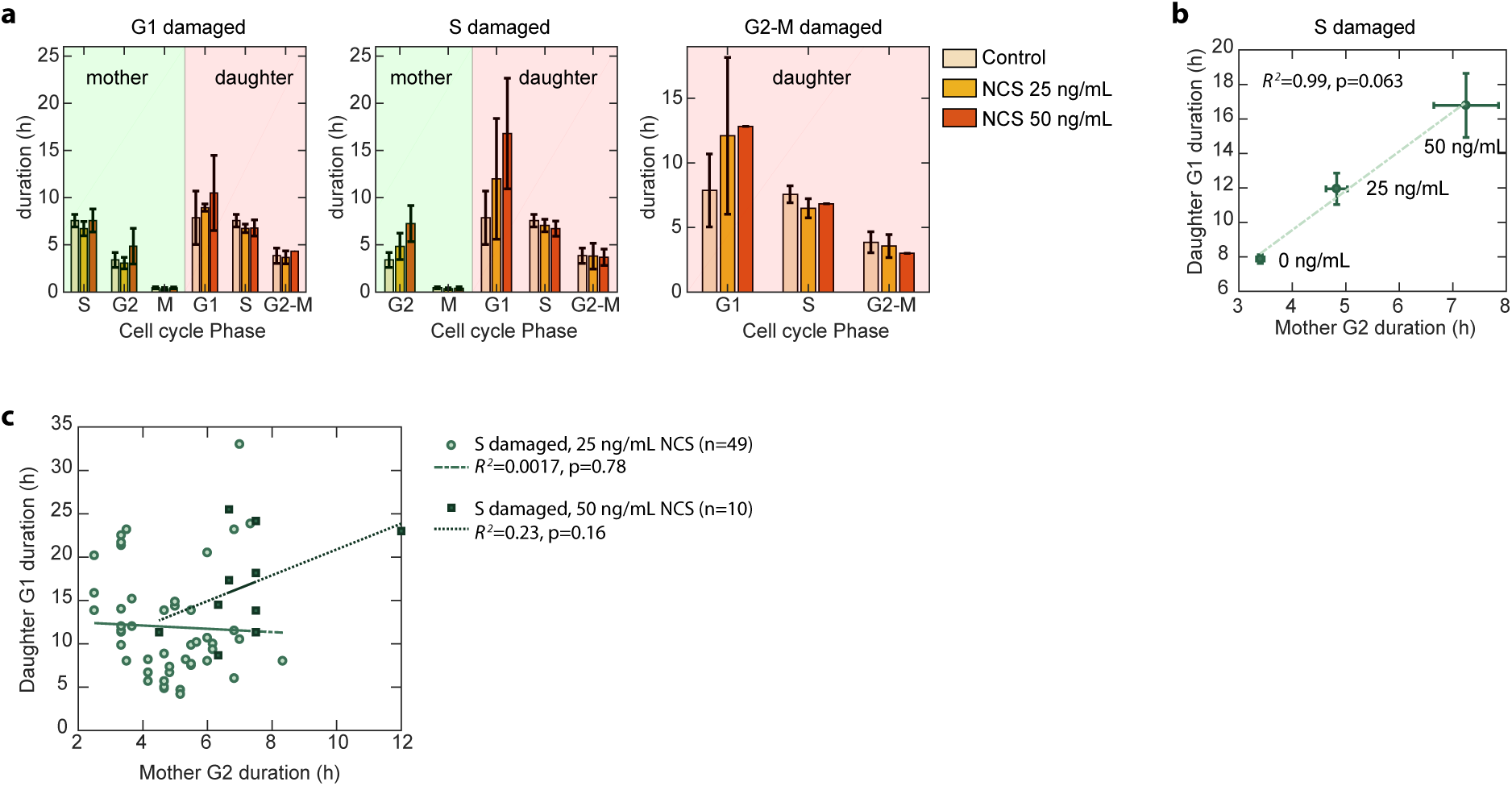
Phase coupling between mother and daughter cells with DNA damage perturbation. **a,** Cell cycle phase durations of RPE cells whose mother cells were treated with NCS during G1 (left panel), S (middle panel), or G2-M (right panel) phases. The cell cycle phase durations of the subsequent cell cycle of the daughter cells were also measured. Error bar represents standard deviation. **b,** Correlation between the population mean G2 duration of the mother cell treated in S phase and the mean G1 durations of their daughter cells, grouped by different NCS concentrations, fitted with linear regression. Error bars represent standard error of mean. **c,** Correlation between the G2 durations of the mother cell treated in S phase and the G1 durations of the daughter cells, fitted with linear regression. Legend indicates the phase in which the mother cell was damaged at the indicated NCS concentrations.

**Figure S6.**
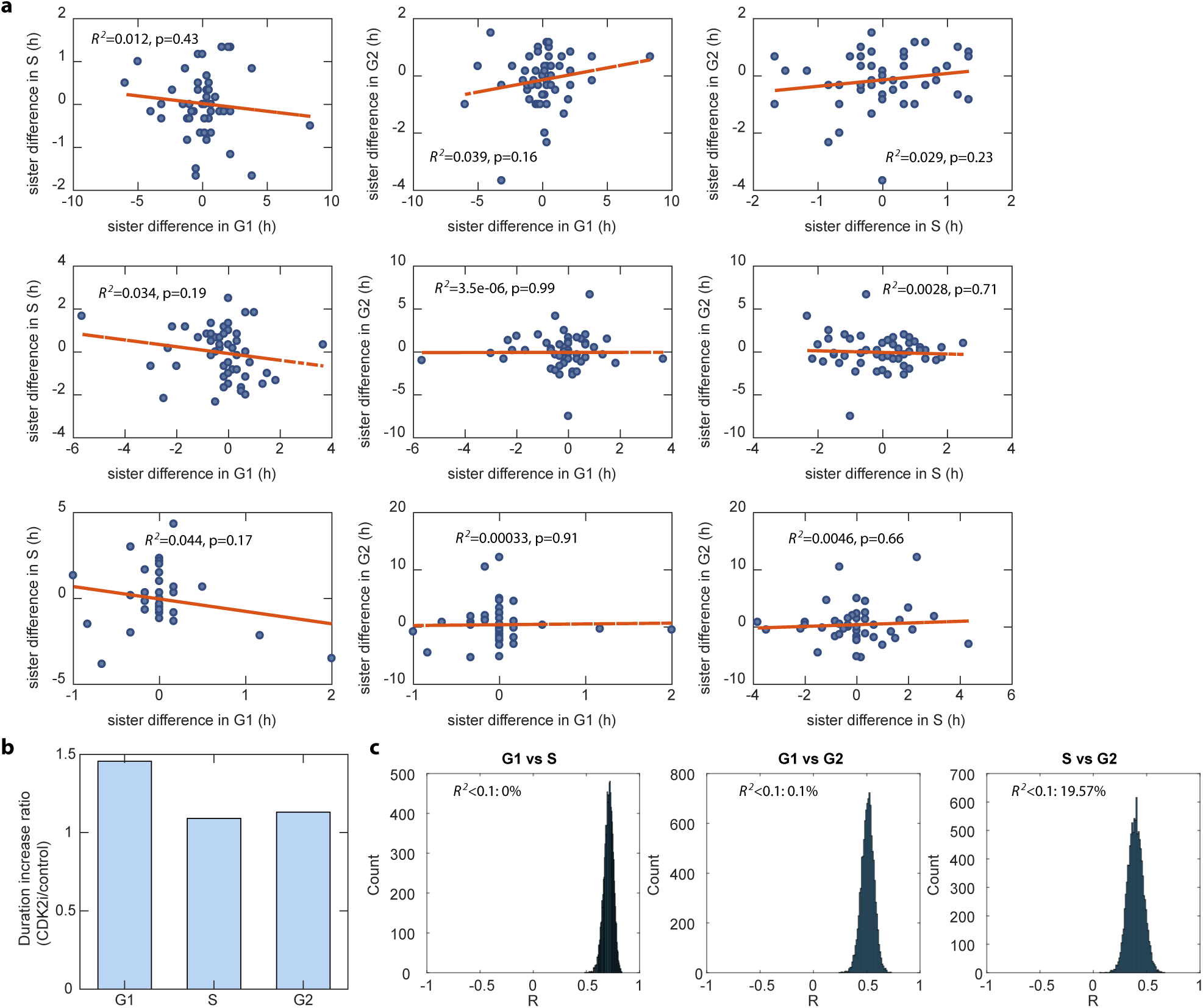
A model for the heritable factors governing phase progression rate. **a,** Correlation plots in the difference in cell cycle phase durations between the sibling cells. To calculate the difference, the subtrahend and minuend were randomly choosen between the sibling cells. n>98. **b,** Ratio of cell cycle phase duration increase with CDK2 inhibitor treatment relative to control. **c,** Non-parametric bootstrap of the distribution of correlation coefficient (R) for RPE under CDK2 inhibitor treatment. n=10,000.

**Figure S7.**
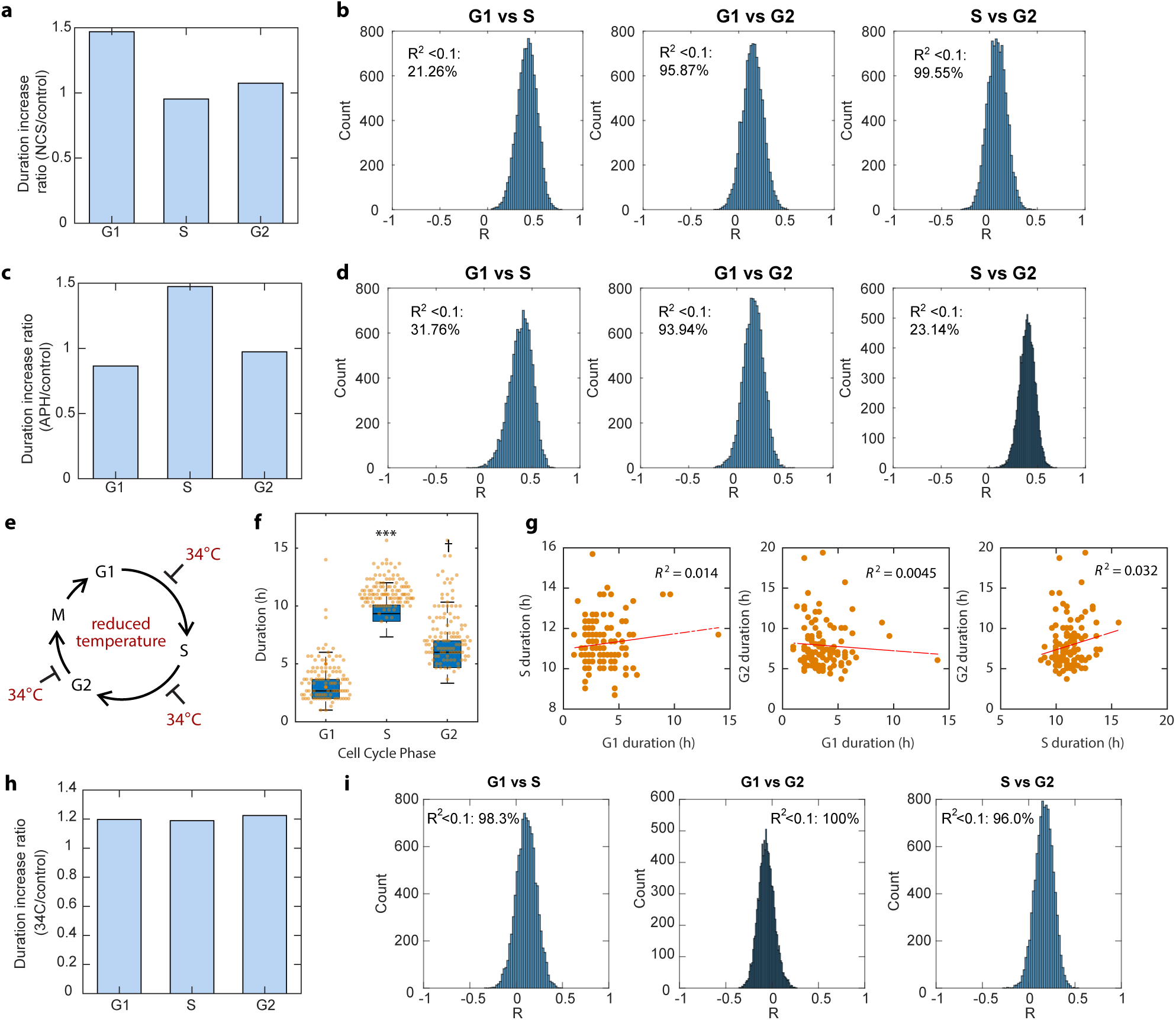
Bootstrap analysis for correlation coefficients between cell cycle phases under perturbation. **a,** Ratio of U2OS’s cell cycle phase duration increase under 100 ng/mL NCS treatment relative to control. **b,** Non-parametric bootstrap of the distribution of correlation coefficient (R) for U2OS under 100 ng/mL NCS treatment. n=10,000. **c,** Ratio of U2OS’s cell cycle phase duration increase under 50 ng/mL APH treatment relative to control. **d,** Non-parametric bootstrap of the distribution of correlation coefficient (R) for U2OS under 50 ng/mL APH treatment. n=10,000. **e,** Schematic of prolonging all phases by growing under 34°C condition. **f,** Cell cycle phase durations of U2OS cells growing under 34°C condition. Boxplots for untreated cells are underlaid for comparison. ***, *P* < 1 × 10^−20^, †, *P* < 1 × 10^−4^, 2-sided Kolmogorov–Smirnov test. *n* = 112. **g,** Pairwise correlation between cell cycle phase durations growing under 34°C condition. **h,** Ratio of U2OS’s cell cycle phase duration increase under 34°C growing condition relative to control. **i,** Non-parametric bootstrap of the distribution of correlation coefficient (R) for U2OS under 34°C growing condition. *n* = 10,000.

## Method Details - Model Description

### 1. Cell cycle progression model simulations and parameter fitting

#### Fitting with the simple Markovian model with a single rate parameter

All simulations and parameter fitting were performed using MATLAB. The durations of cell cycle phase —G1, S, G2, and M —under basal conditions were together fitted to four Erlang distributions with the same rate (*λ*) parameter. The shape (*k*) parameters were restricted to positive integer and were allowed to vary for each cell cycle phase (Figure S3a). The fitting was performed by maximizing the likelihood of observing the experimental data using the *fminsearch* function in MATLAB.

Under the Erlang distribution, the probability of observing a cell of a particular cell cycle phase, for example, G1’s, duration *x*, *f* (*x*; *k, λ*) is

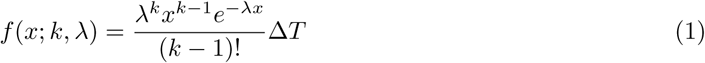

where Δ*T* is the measurement interval.

Then the probability of observing a cell of four cell cycle phase durations x, *f* (x; k*, λ*) is

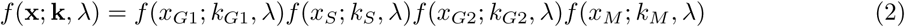

The shape and rate parameters were determined by solving for the maximal likelihood of observing the experimental data:

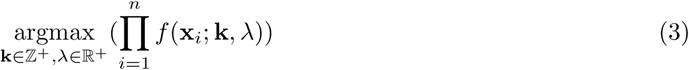

where x*_i_* is the *i*th cell in the experimental data, and *n* is the total number of observed cells.

#### Fitting with the Erlang model with flexible rate parameter

The durations of each cell cycle phase —G1, S, G2, or M —under basal conditions were independently fitted to an Erlang distribution (**Figure 2a**). For each cell cycle phase, we fit the experimental distribution of cell cycle phase durations to obtain the shape (*k*) parameter and the rate (*λ*). For each cell cycle phase, the shape and rate parameters were independently determined by solving for the maximal likelihood of observing the experimental data of each phase:

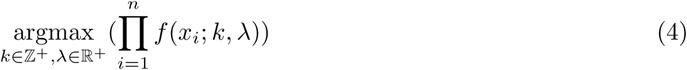

where *x_i_* is the *i*th cell in the experimental data, and *n* is the total number of observed cells.

#### Simulation of cell cycle phase transition

After the fitting with the Erlang model, we obtained 2 parameters for each cell cycle phase and each cell line. Using the estimated parameters, we simulated the progression of cell cycle phase using the Gillespie stochastic algorithm (Figure S3b). Alternatively, because the Erlang distribution is a special case of the Gamma distribution with integer scale parameter, we can generate the phase durations from a gamma distribution in MATLAB (Figure S2b-d):

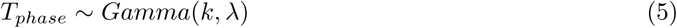

For the normal distribution model, parameters for each cell cycle phase were independently chosen according to the mean (*µ*) and variance (*σ*^2^) of the experimental cell cycle phase durations distributions. The cell cycle phase durations were then simulated from a Gaussian distribution (Figure S2d).

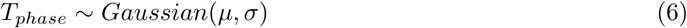

#### Erlang distribution as an approximation of the hypoexponential distribution

Our Erlang model describe the cell cycle phase progression as a series of sub-phase transitions with the same rate *λ*. The relevant biological interpretation of the Erlang model is that each cell cycle phase can be viewed as a multistep biochemical process that needs to be completed sequentially in order to advance to the next cell cycle phase. Biologically, the rate of each sub-phase transition could be different from one another. A model that can account for this flexibility is the hypoexponential distribution, or the generalized Erlang distribution, which allows the rate parameter of each transition to be different. However, the Welch-Satterthwaite equation provides a good approximation of the generic sum of multiple Erlang distributions as one Erlang distribution^1,2^:

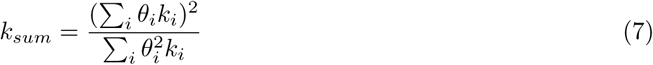

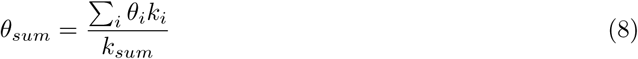

Where the *k_i_* and *θ_i_*are the shape and scale parameters for the *i*th individual Erlang distribution, and the sum of *i* Erlang distributions can be approximated by an Erlang distribution with only two parameters: Erlang(*k_sum_, θ_sum_*)

### 2. The many-for-all model of heritable factors governing cell cycle progression rate

#### Many-for-all model with only phase-coupling factors

The many-for-all model for heritable factors assumes that there are physical factors, called “phase-length factor”, inside the cells that control the rate of cell cycle phase progression. In addition, the levels of these factors can fluctuate throughout the cell cycle but are evenly distributed among sibling cells during mitosis so that sibling cells share similar amounts of the heritable factor. Each type of phase-length factor has shared control among two or more cell cycle phases, exerting an effect (*a*) on multiple cell cycle phase durations by influencing the rates of cell cycle phase progression. The magnitude of the factor effect is proportional to the amount of factor (copy number of molecules). Take G1 and S phase for example, the rate of G1 progression is dependent on the sum of effects among every factor:

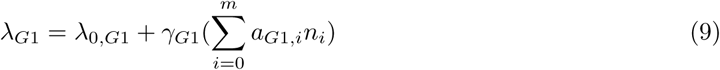

where *λ*_0,*G*1_ is the average progression rate of G1, *γ* is the fraction of progression rate subjected to the control of phase-length factors, *a_G_*_1,*i*_ is the effect coefficient of factor type *i*, *n_i_* is the copy number of factor type *i*, and *m* is the total number of different factor types. The effect coefficients were assumed to follow a normal distribution with mean zero:

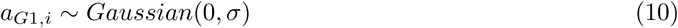

Similarly for S phase:

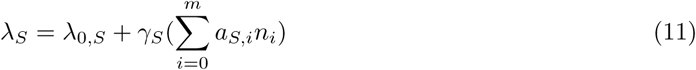

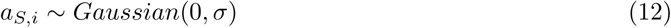

*σ* was chosen to be 0.01 to generate cell cycle phase distributions that resembled experimental data. The copy numbers of each factor type (*n_i_*) for each cell were assumed to follow a Poisson distribution^3,4^, with mean abundance following a lognormal distribution of *µ* = 1000 and *σ* = 0.6^5–7^.

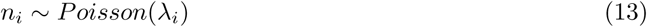

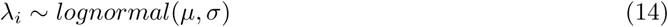

Modeling the factor copy number with a normal distribution with variance equals the mean did not affect the results (data not shown):

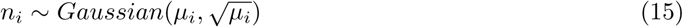

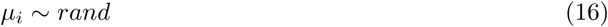

Hence, the G1 duration is

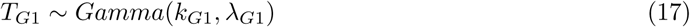

Similarly for S phase:

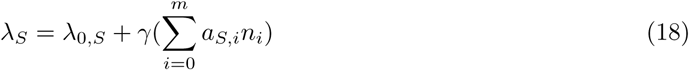

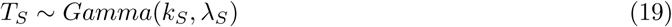

The Pearson correlation coefficients was then calculated by generating 200 cells with G1 and S phase durations using the simulation framework described above (**Figure 4b**).

#### Many-for-all model with both phase-coupling factors and phase-specific factors

In addition to the phase-coupling factors, which has shared control among two or more cell cycle phases, we took into account the presence of phase-specific factors, which affect only one specific cell cycle phase. Take G1 and S phase for example, for G1-specific factors, *a_S,i_*= 0. For S-specific factors, *a_G_*_1_*_,i_* = 0. The rate of G1 progression is dependent on the sum of effects among every factor, including both the phase-coupling factors and the phase-specific factors.

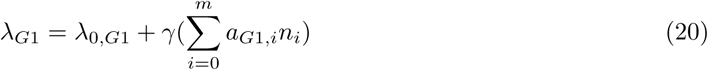

where *λ*_0_*_,G_*_1_ is the average progression rate of G1, *γ* is the fraction of progression rate subjected to the control of phase-length factors, *a_G_*1*_,i_* is the effect coefficient of factor type *i*, *n_i_* is the copy number of factor type *i*, and *m* is the total number of different factor types. The effect coefficients were assumed to follow a normal distribution with mean zero:

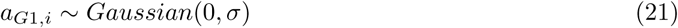

for phase-coupling factors, and equals zero for S phase-specific factors.

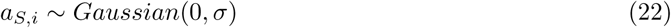

for phase-coupling factors, and equals zero for G1 phase-specific factors.

*σ* was chosen to be 0.01 to generate cell cycle phase distributions that resembled experimental data. The copy numbers of each factor type (*n_i_*) for each cell were assumed to follow a Poisson distribution^3,4^, with mean abundance following a lognormal distribution of *µ* = 1000 and *σ* = 0.6^5–7^.

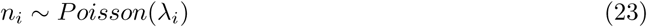

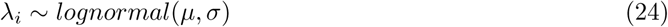

Modeling the factor copy number with a normal distribution with variance equals to mean did not affect the results (data not shown).

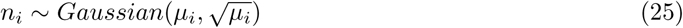

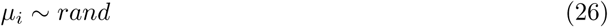

Hence, the G1 duration is

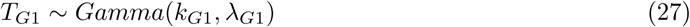

Similarly for S phase:

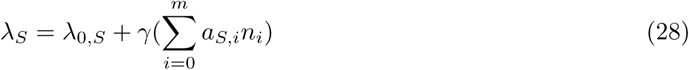

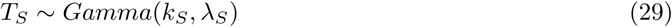

The Pearson correlation coefficients was then calculated by generating 200 cells with G1 and S phase durations using the simulation framework described above (**Figure 4c**).

#### Perturbation of a single phase-coupling factor

The effect of perturbing a single factor was modeled by choosing the type of phase-coupling factor that had the largest product of effect coefficients on two phases. That is, find *i* that maximizes (*a_G_*_1_*_,i_ ×a_S,i_*).

After the *i* was determined, the abundance of that factor was increased by 10 folds. That is, 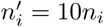. The cell cycle phase durations were simulated similarly as above, except for the increased value of *n_i_* calculated above (**Figure 4d-e**).

#### Requirement of independent assortment of heritable factor into daughter cells

For sibling cells, the factor abundance is assumed to be strongly correlated. That is, the correlation coefficient between the copy number for each factor type i (*ρ_n_*_1*i*,*n*2*i*_) is large. *n*1*_i_* and *n*2*_i_* represent the copy numbers of factor type i for the two sibling cells.

Thus, the difference in cell cycle phase duration between the two sibling cells can be expressed as a function of:

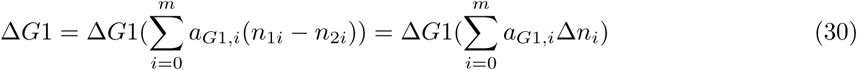

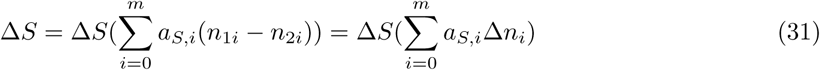

If the segregation of each factor during cell division is not independently distributed, but correlated. That is, if 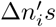 are correlated, then we can rewrite

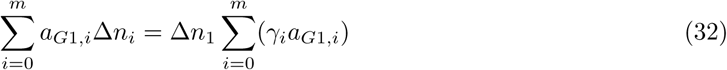

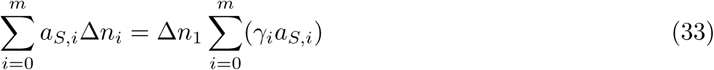

where *γ_i_* is the proportionality terms between 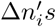 plus the noise term. Under this condition, Δ*G*1 and Δ*S* would be correlated. Our observation that there is no correlation in the differences in cell cycle phase durations between sibling cells, suggesting that the propagation of factors into daughter cells is not interdependent.

